# Mechanical torque promotes bipolarity of the mitotic spindle through multi-centrosomal clustering

**DOI:** 10.1101/2021.11.17.469054

**Authors:** Christopher E. Miles, Jie Zhu, Alex Mogilner

## Abstract

Intracellular forces shape cellular organization and function. One example is the mitotic spindle, a cellular machine consisting of multiple chromosomes and centrosomes which interact via dynamic microtubule filaments and motor proteins, resulting in complicated spatially dependent forces. For a cell to divide properly, is important for the spindle to be bipolar, with chromosomes at the center and multiple centrosomes clustered into two ‘poles’ at opposite sides of the chromosomes. Experimental observations show that in unhealthy cells, the spindle can take on a variety of patterns. What forces drive each of these patterns? It is known that attraction between centrosomes is key to bipolarity, but what the prevents the centrosomes from collapsing into a monopolar configuration? Here, we explore the hypothesis that torque rotating chromosome arms into orientations perpendicular to the centrosome-centromere vector promotes spindle bipolarity. To test this hypothesis, we construct a pairwise-interaction model of the spindle. On a continuum version of the model, an integro-PDE system, we perform linear stability analysis and construct numerical solutions which display a variety of spatial patterns. We also simulate a discrete particle model resulting in a phase diagram that confirms that the spindle bipolarity emerges most robustly with torque. Altogether, our results suggest that rotational forces may play an important role in dictating spindle patterning.

## 1 Introduction

Spatial organization in the interior of cells is intimately linked with cellular function. Consequently, understanding the underlying mechanisms of this organization is a fundamental pursuit in cell biology. One such example is the mitotic spindle, a cellular machine that spatially organizes copied genetic material during cell division. The mitotic spindle has several distinct phases, but here we focus on so-called *metaphase*, where in healthy cells, centrosomes (CSs) are at two opposite ‘poles’ and chromosomes (CHs) aggregated in the middle, on the ‘equator’, into the so called metaphase plate [DM09], as seen in **Fig. 1**. This architecture is crucial for proper segregation of CHs when the cell divides.

**Figure 1:**
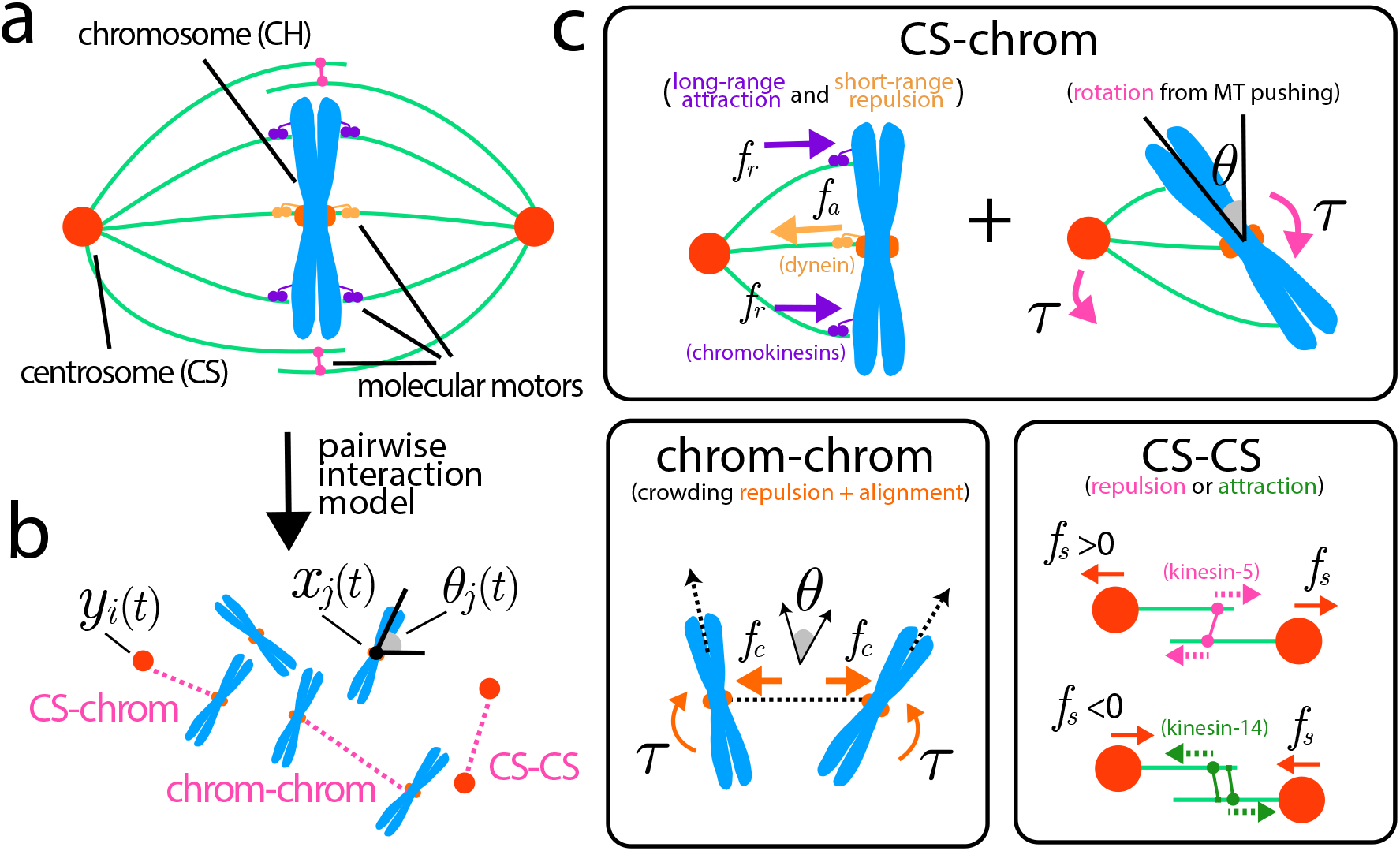
a: The bipolar, mitotic spindle. Centrosomes (CSs) are positioned on opposite sides of a group of chromosomes (CHs). Their interaction is through microtubule filaments and molecular motors associated with them. b: Individual interactions in the model. CS-chrom interactions consist of shortrange repulsion from poleward ejection forces of strength *f*_*r*_ and long-range attraction from motors of strength *f*_*a*_. A resulting torque from these forces which promotes CH being orthogonal to their interaction with the CS (*θ* = 0). CSs attract or repulse each other depending on model parameter *f*_*s*_. CHs repulse each other with strength *f*_*c*_ and align via a local torque.

The spindle is not always bipolar. Various abnormalities in cancer cells (but also some healthy cells) result in the appearance of more than two CSs per cell [RG17]. In these cells, the resulting spindle pattern varies, and can be monopolar, bipolar, or multipolar. [FVM11]. If the resulting pattern is *multipolar*, consisting of CSs aggregating in more than two groups, proper mitosis fails [Bas+08] and these cells often die or display developmental defects [ORA12]. However, cancer cells with multiple CSs can ‘cluster’ these extra components into an apparent bipolar spindle and these cells divide normally [Kwo+08]. Because this clustering allows many cancer cells to proliferate, understanding of the underlying mechanics of spindle patterns is important for development of anti-cancer therapies [ORA12].

Both chemical and mechanical interactions are known to be involved in the formation of the spindle [DM09], but we will focus on purely mechanical in this work. The mechanical forces between the CSs and CHs are primarily generated by microtubules (MTs) and molecular motors, as seen in **Fig. 1**. MTs are anchored in the CSs with their minus ends, while the plus ends grow outward, in random directions and with complex, stochastic dynamics [DM09]. Some of antiparallel MT pairs from two CSs overlap, and a host of motors at the overlaps exert forces on the MTs. Some of these motors (i.e. kinesin-14 [DM09]) generate forces by attaching to one MT with their cargo domains and using the motor head to walk toward the minus end of the other MT, effectively sliding the other MT inward (**Fig. 1C**) and generating the attraction between the respective pair of the CSs. Other motors (i.e. kinesin-5 [Kwo+08]) are bipolar with motor heads on both ends; on the antiparallel MT overlaps these motors walk to the plus ends of both MTs effectively sliding the MTs apart and giving rise to the repulsion between the two CSs (**Fig. 1C**). Yet other motors, dyneins, ‘reel in’ MTs that connect at their plus ends with kinetochores: large protein complexes in the middle of the CHs, resulting in effective attraction between any pair of CS and CH [HS90] (**Fig. 1A,C**). Dynein motors play many roles in mitosis depending on differential localization and regulation. Henceforth, when referencing dynein, we will specify their localization and respective action. However, other MTs run with their plus ends into the chromosome arms, and motors on the arms (chromokinesins), by walking to the MT plus ends, tend to push the MT tips away generating an effective repulsion between a pair of CS and CH, called the polar ejection force [KCH09]. In addition to the motor-generated forces, MT dynamics lead to both pushing (growing MTs pressing on the CH arms) and pulling (shortening MTs effectively pulling the kinetochores hanging on to the disassembling MT ends) forces; mathematically, these forces can be lumped together with the motor-generated forces. (**Fig. 1A,C**) Although this list neglects several important factors (e.g. interaction with the cell boundary/cortex), previous studies have shown these to be sufficient in explaining the origin of of the spindle architecture [Fer+09; Néd02].

However, what happens when the CS number is greater than 2? In this work, we explore this setup through mathematical modeling. Inhibition of the kinesin-14 motor, likely responsible for the CS-CS attraction promotes the frequency of multipolarity [Kwo+08; Bas+08], suggesting that the mutual inter-CS attraction key to the bipolarity [Kwo+08]. However, the simple question arises: why does this CS-CS attraction *not* just aggregate all CSs into just one cluster making the spindle monopolar? Such monopolar spindles were observed in the situations when motors responsible for inter-CS repulsion were inhibited [Kap+00; Fer+09]. One theoretical solution to this problem was proposed in [Cha+20]: attraction of the CSs to the cell cortex at the opposite cell ends can keep two groups of mutually attractive CSs apart. In this study, we wish to find whether such spindle interactions with the cell cortex are necessary, or the multi-CS spindle could remain bipolar autonomously, even without interacting with the cell boundary.

To this end, we explore the idea that a *torque* between the CSs and CHs could complement the forces to ensure the bipolarity of the multi-CS spindle. The idea is as follows: in previous models, CHs were usually considered as material points [Fer+09; Cha+20]. However, they are more accurately represented as double rods ‘glued’ together at the centromere, in the middle, with ‘rabbit ears’ of CH arms stretching from the centromeric region. Mathematically, therefore, the CHs can be described as elongated, oriented objects characterized, besides the coordinates of the center-of-mass, by their orientation angles. This raises the question: can interaction of the CH-CS pair depend not only on their mutual distance, but also on the angle between the CS-CH vector and CH axis? We posit that it can: if the angle deviates from 90 degrees, then one of the CH arms is closer to the CS, and the polar ejection force from the CS onto this proximal arm is greater than that between the CS and another, distal arm. This effect would create a torque that tends to pivot the CS and CH until the CS-CH vector and CH axis are normal to each other (**Fig. 1C**). We hypothesize that such action assists spindle bipolarity, see **Fig. 2**: if the CHs cluster together and align with each other (as is observed in the metaphase plate [HS90; FCR02]), then multiple CSs would be pushed away from most of the space into two sectors that are perpendicular to the CH orientation. This effect would prevent aggregation of the CS groups from two different sides of the CH cluster and lead to eventual clustering of the CSs into just two groups. In order to test this intuition and to determine what spindle states emerge from all these complex interactions, we resort to mathematical modeling.

**Figure 2:**
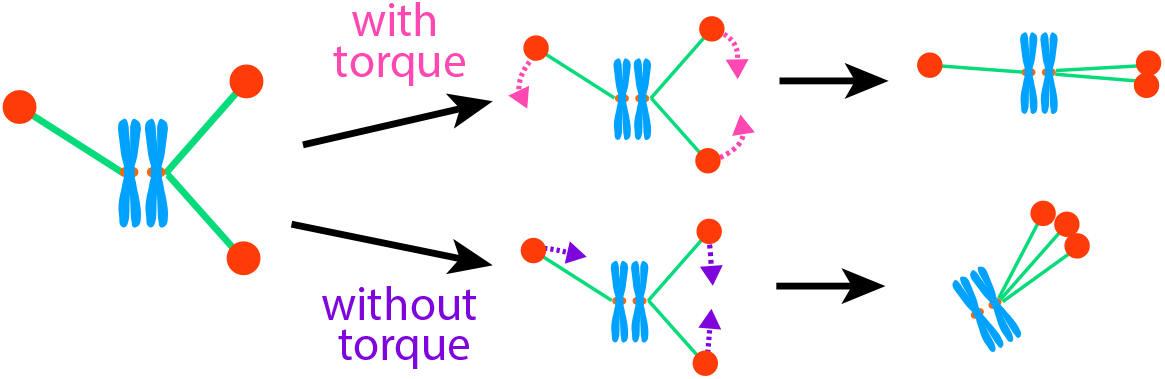
Cartoon explaining intuition behind torque promoting a bipolar spindle. Without torque, centrosomes (CSs) attract each other and aggregate into a monopolar configuration. With torque, CS aggregates still form but are offset by the rotational motion that promoting a bipolar configuration.

Modeling has a long history of helping experiment to elucidate the spindle dynamics ([Arm+15; ZG15; PT16; ZG15; Red+19; Ede+20]). Specifically, force-balance models have been used to probe the spindle structures in one-dimensional (1D) [Néd02; Fer+09], two-dimensional (2D) [ZG15], and in realistic three-dimensional (3d) [Ede+20; Cha+20] geometry. Several types of mathematical models of the spindle have been proposed. There are the most detailed agent-based models (discrete and stochastic) that explicitly simulate biochemical details of individual molecular motors and dynamic elastic MTs [Let+19]. The advantage of such detailed models is unambiguous mapping onto experimentally observed architectures, but the drawback is difficulty of exploring parameter space of the models. In principle, one can roughly average the action of multiple motors and MTs; the resulting mean-field approximation leads to the second type of models: pairwise-‘interacting particle’ models [Fer+09; Man+18], in which the CSs and CHs are modeled as particles driven by interactions through distance-dependent forces. The forces in the particle models often have corresponding potential energy, and so the third type of models - minimizing the total mechanical energy of the spindle - can give us the spindle’s mechanical equilibrium without simulating the transient dynamics [Cha+20]. Lastly, an additional approximation can be made by representing the CSs and CHs as continuous density, rather than as discrete particles. In this study, we avoid the complexity of the most detailed agent-based models and explore the role of the CS-CH torque in the bipolarity of the multi-CS spindles by using the interacting particle and continuous models.

## 2 Continuous and discrete models

### 2.1 Discrete (interacting particles) model

In the model, we consider *i* = 1, …, *N*_*S*_ CSs with positions *y*_*i*_(*t*) ∈ ℝ^2^ and *j* = 1, …, *N*_*C*_ CHs with positions *x*_*j*_(*t*) ∈ ℝ^2^ and orientations *θ*_*j*_(*t*). Each CS (and each CH) interacts with all other CSs and all CHs (respectively, all CSs) by pairwise, distance-dependent interactions, see **Fig. 1**. The evolution of all positions and orientations is described by the equations

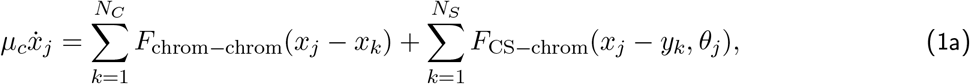

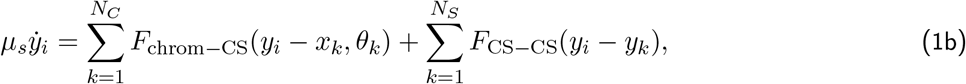

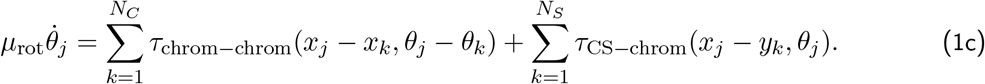

Here, as is conventional in cellular system, we use the overdamped mechanical equations, in which linear and angular velocities are proportional to forces and torques, respectively, and inertial terms are negligible due to the low Reynolds numbers. Parameters *μ*_*c*_, *μ*_*s*_, *μ*_rot_ are the effective viscous drags for the CH, CS and CH arms rotation, respectively. We introduce the distance dependence of the forces and torques and model parameters below. The CS and CH coordinates are the 2D vectors; 0 < *θ*_*j*_(*t*) < *π*. Numerical integration of the first order ODEs of the discrete model is straightforward and standard.

### 2.2 Continuous model

As is common in pairwise interacting particle models, we seek qualitative insight of our model by considering the limit of dense particles, or the so-called *mean-field* limit. Citing rigorous arguments such as [BV05; CCH14] but omitting further detail here, we introduce densities of CSs (*S*(*x, t*)) and CHs (*C*(*x, θ, t*)). These densities arise in the limit of dense particles, 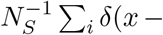 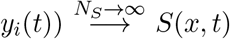 and 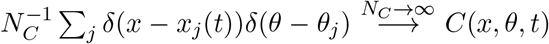. In this limit, integrals become sums, and the resulting continuous system of integro-PDEs has the form

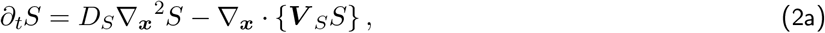

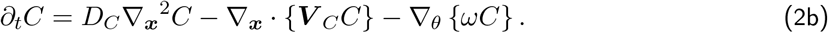

In the continuous system we add diffusion terms to the linear motion of CSs and CHs that represent random movements of the particles resulting from stochastic perturbations not present in (1) but could be included. *D*_*S*_ and *D*_*C*_ are the respective diffusion coefficients. Note that we do not add the rotational diffusion, and we also consider initial conditions such that 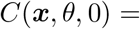 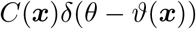. Therefore, the hyperbolic character of the PDEs with respect to the angular variable, ensures the angular distribution remains such that there is a deterministic orientation angle at each spatial point. Although it is unclear a priori if the mean-field limit is appropriate for our system of study with a finite number of interacting bodies, we will later compare qualitative results of this analysis with explicit simulations of the particle system (1). The linear and angular velocities in the transport equations are given by the convolution integrals:

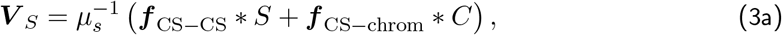

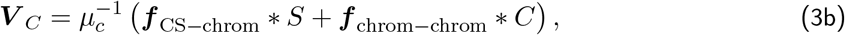

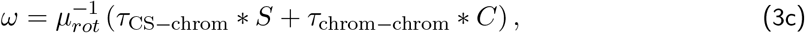

where the convolution operation is defined by

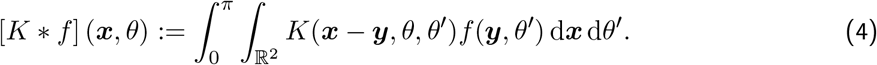

We describe the distance dependence of the forces and torques and model parameters below.

## 3 Results

The continuous integro-PDE model of the spindle is mathematically similar to spatial models of animal grouping and swarming. The latter have a rich history of study [MEK99; BT11; BT13; CCH14; SRS15], including stability, bifurcations, and numerical analyses, or inferring interactions directly [LLEK10; Lu+19]. Here, we focus on characterizing the qualitative behavior of equilibria. following along the lines of significant recent progress found in. We utilize tools developed for the animal grouping models to study the spindle model. The choices of the parameter values in the model are explained and justified below.

### 3.1 Linear stability analysis in 1D

In order to isolate the effect of torque on the continuous model, we consider a much simpler setup: the continuous model in one spatial dimension (1D) with no boundaries and no alignment or torque terms. In this case, we use the forces exponentially decreasing with the distance between pairs of interacting particles:

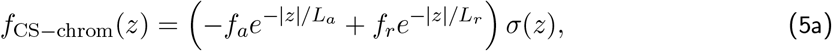

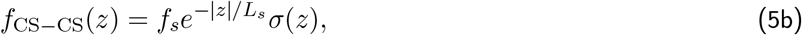

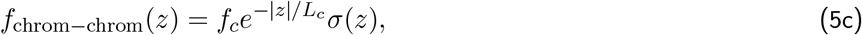

where *σ*(*z*) = sign(*z*). Here *f*_*a*_ and *f*_*r*_ (both are positive parameters) are the amplitudes of the attraction (pulling the centromere of the CH toward the CS) and repulsion (polar ejection force: pushing the CH arms away from the CS) forces; *L*_*a*_ and *L*_*r*_ are the spatial ranges of the respective forces, all chosen to satisfy the biologically relevant regime of H-stability [D’O+06]. The parameters *f*_*s*_ and *f*_*c*_ are the amplitudes of interactions between the CS pairs and CH pairs, respectively. There is a short-distance steric repulsion between the CH pairs, so *f*_*c*_ > 0, however, the CSs can either repel (if *f*_*s*_ > 0) or attract (if *f*_*s*_ < 0) each other. *L*_*s*_ and *L*_*c*_ are the spatial ranges of the respective forces. Three of the force amplitudes (*f*_*s*_, *f*_*a*_ and *f*_*r*_) are proportional to respective net motor forces generated at the interpolar MT overlaps by kinesin-14 and −5 and possibly cytoplasmic dynein, at the kinetochores by dynein, and at the CH arms by chromokinesins, respectively. These amplitudes are proportional to the characteristic force per motor multiplied by the average number of the respective motors. The exponential distance dependence arises from the following factor: the farther apart the interacting organelles are, the smaller number of MT plus ends reach from one organelle to the other (or smaller MT-MT overlap between the organelles). In the simplest case, the MT length distribution is exponential, in which case previous models showed that the forces decrease exponentially with the distance [Fer+09].

In 1D, the system of equations (2) becomes

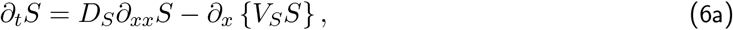

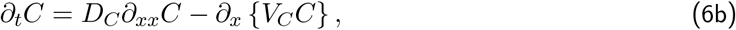

where CS and CH velocities are given by the convolutions of respective forces and densities (3), now take the form

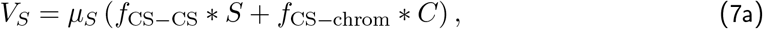

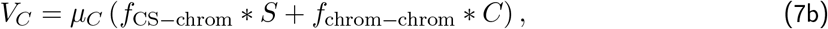

and the convolution (4) becomes

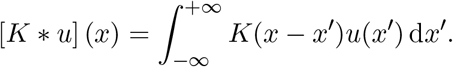

We start the analysis by noting that this 1D model sustains a homogeneous steady-state solution: ***u***(*x*) = [*S*(*x*), *C*(*x*)] = [*S*_0_, *C*_0_]. The system is invariant to translations, so we consider a perturbation from this equilibrium of the form:

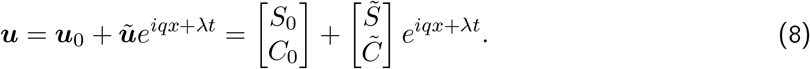

Substituting this perturbation into (6) and keeping only terms that are linear with respect to 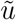, we obtain the linear system of algebraic equations:

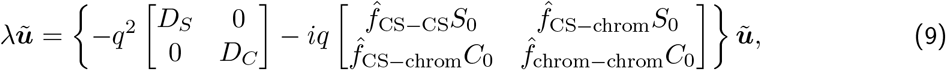

in which the Fourier transforms for a general *f* (*z*) are defined by

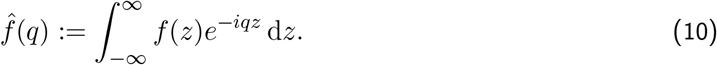

Rewriting these equations in the matrix form:

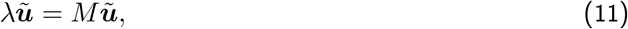

the stability boils down to the eigenvalues of the matrix *M* . This matrix can be computed explicitly for our choice of the inter-particle forces (abbreviating *k* := *q*^2^ > 0):

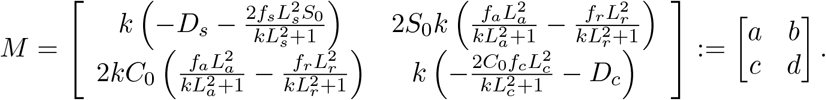

The eigenvalues of this 2 × 2 matrix are real and give us the dispersion relation,

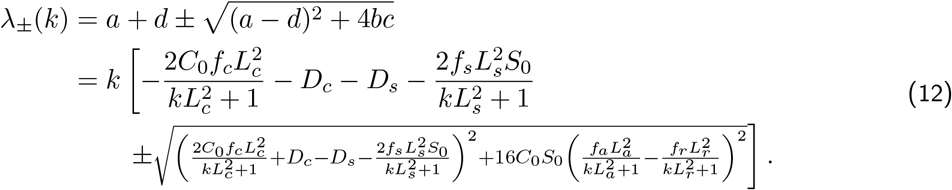

This expression is challenging to understand directly. We can immediately see that *λ*(0) = 0 is expected due to the conservation of the number of particles. Beyond that, since it is always useful to detect instabilities at low wave-numbers, which correspond to the aggregation-type instabilities, we can expand the dispersion relation into Taylor series around *k* = 0:

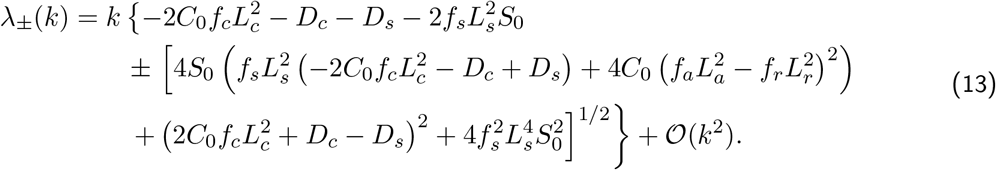

This provides the sufficient (but not necessary) condition for an instability to occur:

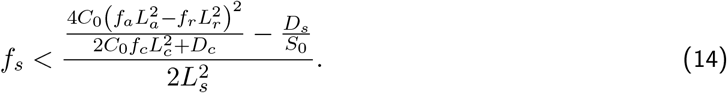

Analysis of this instability criterion indicate that, intuitively, attraction between the CSs (*f*_*s*_ < 0), greater range of the CS-CS interaction, weaker repulsion between the CH pairs and smaller diffusion coefficients promotes the aggregation instability. The quadratic term in the numerator of the aggregation instability criterion indicates that the instability is promoted if net CS-CS attraction (quantified by the product 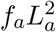) is either much greater or much smaller, than the net CS-CS repulsion (quantified by the product 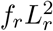).

To gain further insight from the dispersion relation (12) we made the following parameter selection: *D*_*s*_ = 1, *D*_*c*_ = 10, *L*_*a*_ = 10, *L*_*s*_ = 7.5, *L*_*c*_ = 1, *L*_*r*_ = 5, *f*_*a*_ = 1, *f*_*c*_ = 0.5, *S*_0_ = 1, *C*_0_ = 4. The rationale for these choices is as follows. Average CS density is chosen as the unit of density, as there are a few tens of CHs in the spindle, while usually less than 10 CSs [Kwo+08], so we choose the ratio of 4 CHs per 1 CS. The CS diffusion coefficient is set to 1 arbitrarily because there is no respective experimental data and because the qualitative dependence of the results on the values of the diffusion coefficients is relatively trivial. The CH diffusion coefficient is set to 10 because the CHs are likely to be more mobile than the CSs [Fer+09], likely due to CSs moving together with their large MT asters. The ranges of interactions are measured in units of microns. The long-range of attraction between the CSs and centromere regions of the CHs is set to 10, which is on the order of the spindle size [Kwo+08; Fer+09]. The repulsion range between the CSs and chromosome arms is chosen twice shorter than the attraction range to make sure that there is a preferred stable distance between a pair of CS and CH equal to roughly half the spindle length, as observed in metaphase [Fer+09]. The range of interactions between the CS pairs is chosen to be on the same order as those between the CSs and CHs, assuming that the respective MTs have similar lengths. The range of the inter-CH steric repulsion is set to 1, on the order of the CH size. We lump the mobility coefficients together with the amplitude of the forces, so effectively parameters *f* correspond to the velocity amplitudes. We choose the amplitude of the CS-CH attractive velocity to be the velocity unit; the amplitude of the CH-CH repulsive velocity is chosen arbitrarily (as respective biophysics was not investigated before), of the same order of magnitude. Then, we explore the stability as function of two parameters, interaction amplitude between CS pairs *f*_*s*_ and repulsion amplitude between CS and CH arms *f*_*r*_. The results are shown in **Fig. 3.**

**Figure 3:**
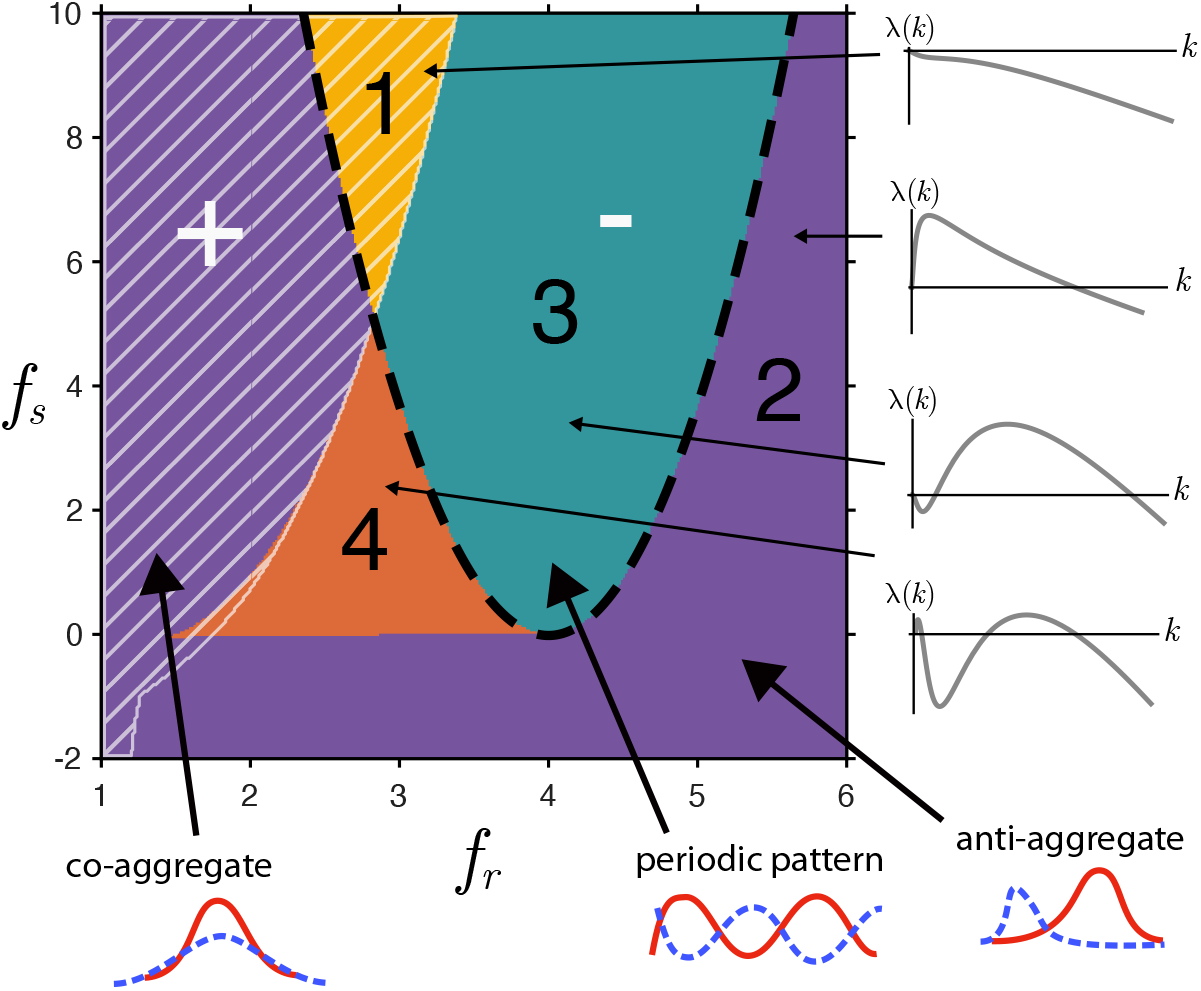
Results of the stability analysis on the 1D model around the spatially homogeneous state. Colors and numbers indicate the number of zeros of the dispersion kernel (12) and black line corresponding to the stability condition in (14) arising the linearization around *k* = 0. The hatched (+) region corresponds to where the eigenvector *u* = [*z,* 1] corresponding to the eigenvalue *λ*(*k*\*), where *k*\* = arg max_*k*_(*λ*) has sign *z* > 0.

From the figure, we see that rich spatial patterning emerges from even the 1D model: there are four different possible stability regimes. The trivial stable equilibrium exists, interestingly, only in a small sliver (region 1) of the parameter space that corresponds to roughly balanced CS-CH attraction/repulsion and to significant inter-CS repulsion. If CS pairs attract (part of region 2 corresponding to negative values of *f*_*s*_), then the CSs aggregate. In order to understand the relative CS-CH localization, we numerically found the eigenvectors corresponding to the eigenvalues of the linear stability equations and deduced the spatial pattern corresponding to the dominant unstable mode close to the equilibrium from relative signs of the harmonics corresponding to the CS and CH densities. That is, we compute the eigenvector *u* = [*z,* 1] corresponding to the eigenvalue compute *λ*(*k*\*), where *k*\* = arg max_*k*_(*λ*) to check the sign of *z*. In **Fig. 3,** the negative sign corresponds to the CS and CH densities in anti-phase (segregated CSs and CHs), while the positive sign means the co-aggregation of the CSs and CHs. We find that when the repulsion between the CSs and CH arms is very weak (left part of region 2), then even if CSs repel each other, the CSs and CHs co-aggregate (plus sign) because inter-species attraction overwhelms intra-species repulsion. On the other hand, when the repulsion between the CSs and CH arms is very strong, (right part of region 2), then when CSs repel each other, the CSs and CHs aggregate (minus sign) into two opposite parts of space (inter-species repulsion overwhelms intra-species repulsion). The most interesting pattern emerges when the CS-CH attraction/repulsion roughly balance each other, while CSs repel each other (region 3). In this region, the periodic pattern of equidistantly grouped interspersed CS and CH clusters emerges: the dominant CS-CH interaction keeps the CS and CH clusters at preferable distance, overwhelming the CS-CS and CH-CH repulsion. Lastly, there is a small sliver of parameter space (region 4), in which the dispersion relation has two maxima, one corresponding to the unstable aggregation mode (small *k*), another - to the periodic spatial instability. The linear stability analysis cannot say which of these patterns will emerge [MEK96], but the linear stability analysis otherwise provides great insight. It is natural to consider whether this stability analysis extends to the 2D model. In the supplementary materials, we follow recent work [FHK11; Kol+11; KHP13; CK14; OHS17; OEK18] to analytically establish existence of ring equilibria of a *modified* model that is qualitatively like the one proposed here. Instead, we turn to numerical simulations of the original model to explore further.

### 3.2 Numerical solutions of the 2D continuous model with torque

We solve the full integro-PDE system (2) and (3) numerically as follows: we discretize the space into 30×30 grid ( Δ*x* ≈ 1 micron), and the angular variable is discretized by 8 equidistant points (Δ*θ* = *π*/8). At each computational step, the convolution integrals are computed by using trapezoidal rule. Then, the advection-diffusion equations are solved by using Crank-Nicolson method. The boundary conditions are no-flux in both spatial dimensions and periodic boundary conditions in the angular direction. The simulations were run for a fixed amount of time, chosen such that an apparent equilibrium was attained. Snapshots of the simulations can be seen in **Fig. 4**. In red, the CS density is shown, and in blue, CH density. Color intensity corresponds to magnitude of the density. Each spatial point is initiated with a single orientation *θ* and the evolution of *θ* is therefore deterministic, and shown in the thin white line. The initial conditions, distance dependence of the forces and parameter values are discussed below.

**Figure 4:**
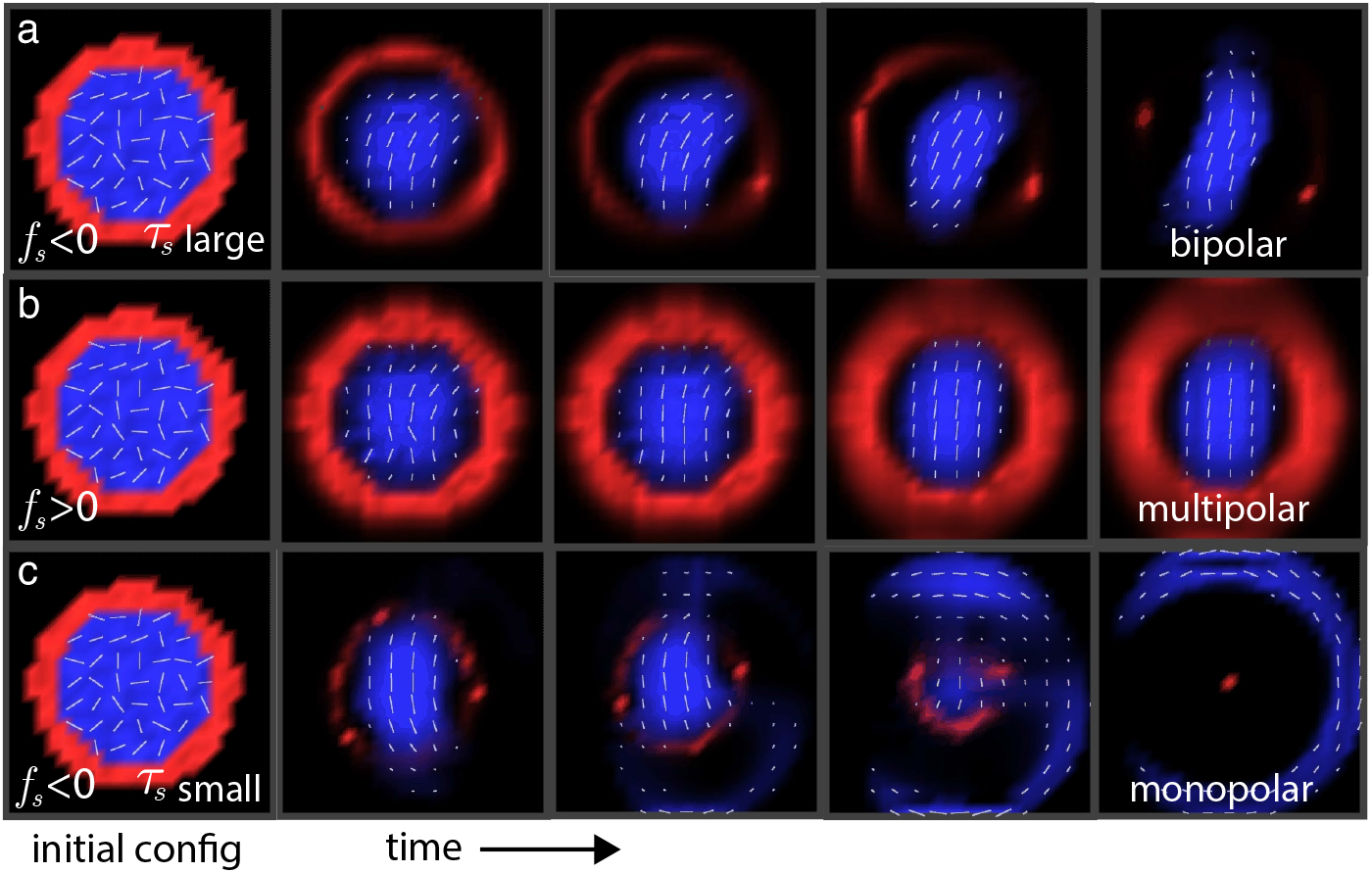
Numerical solutions to the mean-field integro-PDE equations (2) for three parameter sets, each for five time points with identical initial conditions. In red, the centrosomes (CS), and blue, chromosomes (CH). Intensity of color shows magnitude of density and overlaid white lines indicate orientation *θ*. In panel a, the condition with CS-CS attraction (*f*_*s*_ < 0) with strong CS-chrom torque (*τ*_*s*_ large) is shown, resulting in a bipolar spindle. In panel b, CS-CS repulsion (*f*_*s*_ > 0) leads to multipolar spindles. In panel c, when the torque *τ*_*s*_ becomes too small, even with CS-CS attraction (*f*_*s*_ < 0), the spindle becomes monopolar.

We solved the model equations numerically using the numerical methods explained in section 2 and the following distance dependence for the forces:

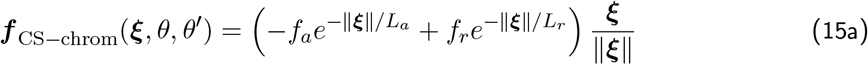

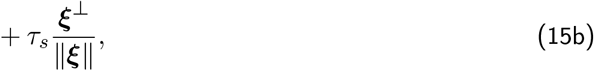

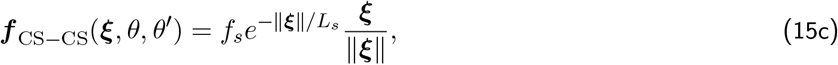

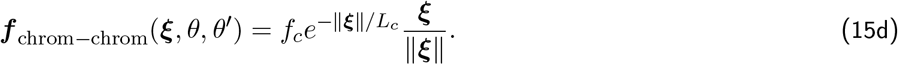

The meaning of the parameters for the forces is the same as that explained for the 1D model. We also use the following torque terms:

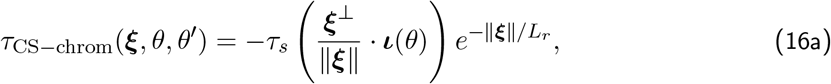

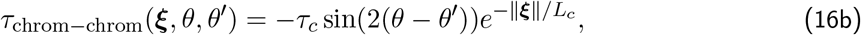

where ***ι*** is the unit-vector in the direction *θ*. The first torque effectively measures the angle between *ι*(*θ*) and ***ξ*** = ***x*** − ***x***′ and evolves toward equilibria of 0 and *π*.

In the simulations, we fixed the majority of the model parameters as follows: *f*_*a*_ = 1, *f*_*c*_ = 1, *f*_*r*_ = 5, *L*_*a*_ = 15, *L*_*c*_ = 0.5, *L*_*r*_ = 5, *L*_*s*_ = 5, in the same units, on the same order of magnitude and based on the same logic as was explained in the 1D model stability analysis. Similarly, the viscous drag coefficients were lumped with the force terms, as explained above. We choose torque amplitudes on the order of unity (characteristic force amplitude on the order of unity multiplied by the CH arm length on the order of 1 micron, which is our length unit). We choose the relatively weak strength of the CH alignment torque, *τ*_*c*_ = 0.2. Lastly, we chose relatively small diffusion coefficients, *D*_*c*_ = 0.2, *D*_*s*_ = 0.2 to provide less influence than the advective terms in lieu of any other knowledge of these parameters.

Then, we varied two parameters: *f*_*s*_, the amplitude of the CS-CS interaction (from −2 to 2), and *τ_s_*, the amplitude of the torque on the CSs (from 0 to 2). As the initial condition, we used the physiologically meaningful CS and CH distributions, as shown in the left column of **Fig. 4**: CHs distributed evenly in a disc and oriented randomly (same as the random distribution of the chromosomes in the nucleus at the onset of mitosis), and CSs distributed uniformly in the ring at the periphery of the CH disc (at prophase, multiple CSs are scattered near the nuclear envelope).

We found that three qualitatively different solutions emerged. For positive values of *f*_*s*_ (CS-CS repulsion), at any value of *τ*_*s*_, the multipolar spindle solutions evolved (**Fig. 4**, middle row, movie 1). Specifically, the CH initial disc-like distribution at the center remained, but the disc radius decreased to an equilibrium (the disc edge smeared due to the diffusion). Meanwhile, the CHs largely align with one another. The ring-like CS distribution also stayed, but interestingly, the ring became non-uniform due to the torque action: the CSs became more concentrated at the ‘poles’, near the direction normal to that of the CH alignment, and depleted from the ‘equator’, near the direction parallel to that of the CH alignment. For negative values of *f*_*s*_ (CS-CS attraction), two different solutions evolved, depending on the torque magnitude. At weak torque (*τ*_*s*_ < 0.5), the monopolar spindle solutions evolved (**Fig. 4**, lower row, movie 2). Transiently, the initial ring of the CSs collapsed into a few clusters, and these clusters, attracting to each other, pushed into the CH disc. Multiple small CS clusters do not constrain the CH density enough in space, and the effective CH diffusion allows the CH density to ‘leak’ outward between the CS clusters. This, in turn, allows the mutual CS-CS attraction to bring the CS clusters closer together, ‘invading’ the CH density and displacing it to the periphery. Eventually, the CSs merged into the single cluster at the center, while the CHs formed the ring (with a break at the side). Locally, the CHs became aligned with the circumference of this ring. The break in the CH ring visible in **Fig. 4**, is a consequence of the initial conditions. This break is not permanent: over a long time (the short-range CH-CH repulsion does not accelerate the ring closure), the effective CH diffusion will spread the CH density evenly in the ring.

Lastly, for negative *f*_*s*_ and strong torque (*τ*_*s*_ > 1.5), the bipolar spindle solutions evolved (**Fig. 4**, upper row, movie 3). The CHs remained in the center but condensed into an ellipse simultaneously aligning along the long axis of the ellipse, resembling the metaphase plate. The CSs remained at the periphery but condensed by torque into two opposite clusters at the spindle poles. At the intermediate torque (0.5 < *τ*_*s*_ < 1.5), monopolar spindles tended to evolve at greater negative values of *f*_*s*_, and bipolar spindles - at smaller negative values of *f*_*s*_. In principle, the parameter space of the continuous model can be fully explored, and respective phase diagram can be sketched. However, the continuous model is merely an approximation to the the discrete model in the limit of a continuous density. As there are only tens of CHs and but a few CSs, whether these lessons hold for these numbers is unclear. Therefore, we resort to the discrete model to confirm their existence even in this scenario.

### 3.3 Spindle configurations in the discrete particle model

We finally turn to numerical simulations of the discrete 2D model of the interacting particles with torque (1). We use the same interaction functions described in the full 2D integro-PDE simulations.

The meaning of the parameters for the forces is again the same as previously mentioned. In the simulations, we kept most of the parameters fixed as follows: *f*_*a*_ = 1, *f*_*c*_ = 1, *f*_*r*_ = 5, *L*_*a*_ = 15, *L*_*c*_ = 1, *L*_*r*_ = 5, *L*_*s*_ = 7.5, in the same units, on the same order of magnitude and based on the same logic as was explained in the 1D model stability analysis. Similarly, the viscous drag coefficients were lumped with the force terms, as explained above. We choose torque amplitudes on the order of unity (characteristic force amplitude on the order of unity multiplied by the CH arm length on the order of 1 micron, which is our length unit). We choose the relatively weak strength of the CH alignment torque, *τ*_*c*_ = 0.5.

We simulate this model with *N*_*s*_ = 10 and *N*_*c*_ = 20 with CHs initially placed uniformly and randomly within the disc ||*x*|| ≤ 10 and CSs initially spaced randomly and uniformly along a circumference with radius ||*x*|| = 10 and the center being the center of the CH disc (movies 4-7). This initial condition corresponds to biologically meaningful initial condition at the prometaphase onset, in which the CHs are at the center, filling the former nuclear sphere after the nuclear envelope breakdown, while the CSs are surrounding the former nuclear envelope. The simulations are terminated when an apparent equilibrium has been reached, defined by the timestep when the displacement for all particles is below 10^−4^ microns.

We then varied two key parameters, *f*_*s*_, *τ*_*s*_, and found that depending on the region of this 2D parameter space, four qualitatively different spindle configurations evolved (**Fig. 5**, movies 4-7). For positive values of *f*_*s*_ (CS-CS repulsion), multipolar spindles evolve (**Fig. 5**, movie 4), with aligned CHs in one cluster at the center, and CSs spread individually and almost equidistantly along the circle around the CHs. In a triangular part of the parameter space with strong CS-CS attraction and weak CS-CH torque, monopolar spindles evolved with all CSs in one cluster at the center, and the CHs distributed almost uniformly along the circle around the CSs and aligned locally along the circumference (**Fig. 5**, movie 5). In most of the rest of the parameter space, corresponding to CS-CS attraction and moderate CS-CH torque, the bipolar spindles evolved with the aligned CH cluster at the center and two CS clusters at the opposite sides of the CHs, so that the CS-CS axis is normal to the CH orientation (**Fig. 5**, movie 6). All these three spindle states and regions of the parameter space corresponding to them predicted by the discrete 2D model are similar to those predicted by the continuous 2D model. Interestingly, the discrete model predicts the fourth possible spindle state (**Fig. 5**, movie 7) not captured by the continuous model. This peculiar state corresponds to a multipolar spindle, but of a special kind: the CSs are not scattered in space individually, but rather grouped into more than two clusters. Furthermore, the CHs aggregate into more than one cluster, in each of which the CHs are aligned with each other, but CH orientation in different clusters is different. These spindle states correspond to large values of CS-CH torque and relatively weak CS-CS attraction. It is easy to understand the origin of these states: when, depending on the initial conditions, such state emerges, due to random initial proximity of sub-groups of the mutually attractive CSs and mutually aligning CHs, then multiple CS clusters do not merge further, because the strong CS-CH torque puts barriers in the way of the CS-CS attraction. Also note that in some regions of the parameter space multiple spindle states emerge at the same parameter values, depending on random initial conditions. These boundary regions are the only locations of parameter space with observable variation between the number of emergent CS clusters.

**Figure 5:**
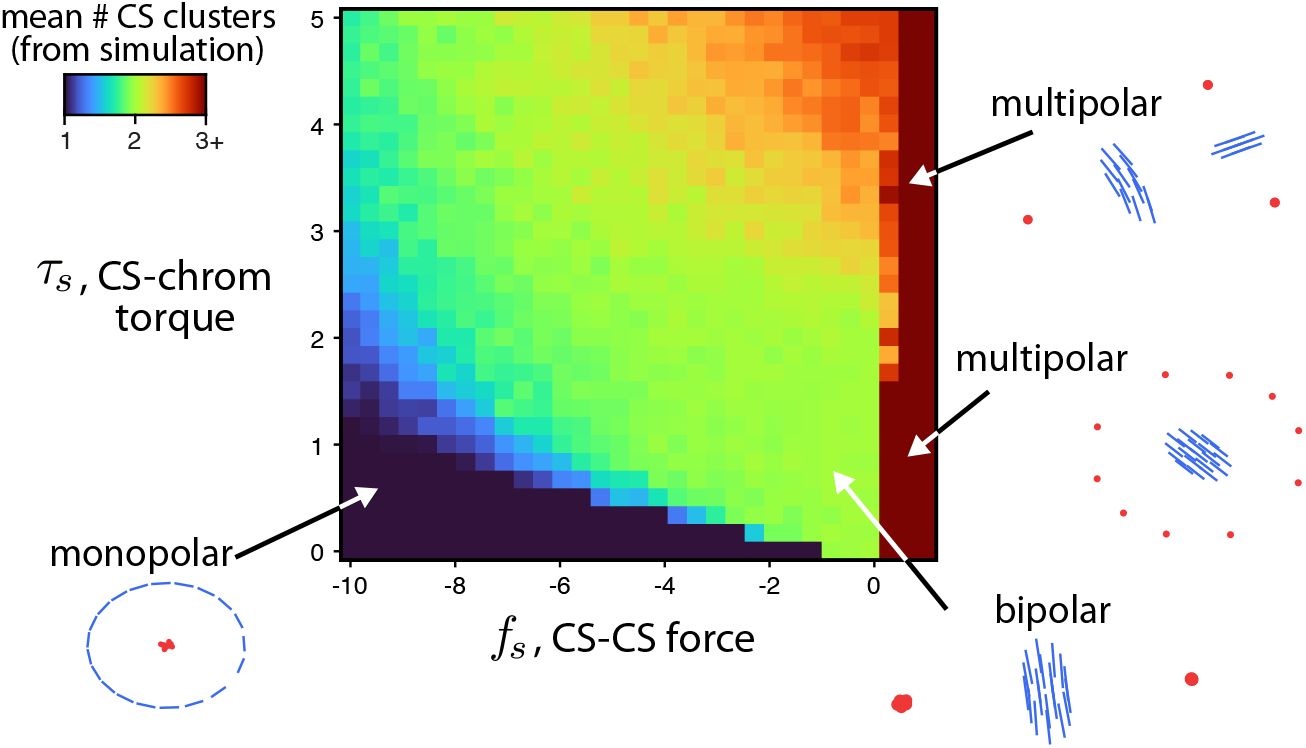
Results of numerical simulations of discrete particle model (1) for varying parameters *f*_*s*_ and *τ*_*s*_. For each set of parameters, 100 simulations with random initial conditions are computed until apparent equilibrium is reached. The mean number of CS clusters over these trials is shown, along with characteristic cartoons of the corresponding typical equilibria. This phase diagram supports the claim that CS-CS attraction *f*_*s*_ < 0 and CS-CH torque *τ*_*s*_ > 0 promote bipolarity.

## 4 Discussion

The stability analysis of the simplified 1D continuous model without torque, expectedly, predicts aggregation of all CSs into a single cluster when CSs mutually attract. Unexpectedly, this model predicts a rich variety of spatial patterns when CSs repel each other, including either co-aggregation or segregation of CSs and CHs and periodic patterns of multiple intermittent CS and CH clusters. The numerical solutions of the 2D continuous model with the CS-CH torque then confirm the hypothesis that the CS-CH torque combined with the CS-CS attraction result in the bipolar spindle configuration. To test whether these predictions survive the transition to the finite discrete system, and to explore the parameter space systematically, we then numerically simulate the discrete model with torque and find that the bipolar spindles indeed evolve if moderate CS-CH torque accompanies limited CS-CS attraction. Notably, greater torque leads to the spindle configurations with not only multiple CS clusters, but also with multiple CH groups.

The modeling predictions qualitatively agree with a few experimental observations: CS clustering is known to be promoted by upregulation of kinesin-14 [Kwo+08] and dynein [Qui+05] motors which generate CS-CS attraction, noting that this interpretation relates specifically to cytoplasmic dynein. CSs that lack CHs between them do not form a stable spindle-like MT array [FCR02], exactly as our models predict. Multiple independent metaphase plates (CH clusters) were also observed [Dun15], similar to the predicted multi-CH-cluster spindles emerging when the torque is too great, as seen in **Fig. 5**. Normally though, multipolar spindles are characterized by a single, interconnected cluster of CHs at the center, with individual or clustered CSs at the periphery [Bau+20]. The observed CH distribution in this single CH cluster within multipolar spindles has branched, Y-, V-, or T-shaped configurations, with CHs aligning between multiple spindle poles [Hen75; WW96; Bau+20; Gou+20], which resembles the model-predicted multipolar state at high torque. (See also the transient Y-shaped CH distribution in movie 7). On the other hand, the predicted multipolar spindle with the CSs encircling the single aligned metaphase CH plate was never observed, which perhaps indicates that the CS-CS force is never repulsive at this mitotic stage. Beyond equilibria, simulations ending in bipolar spindles commonly displayed transient multipolarity, also noted experimentally [SC12]. Lastly, we are not proposing that the CS-CH torque is the only factor preventing the CS-CS attraction from forming monopolar spindles. Indeed, attraction of the CSs to the cell boundary also supports the bipolarity by competing with the CS-CS attraction and separating two CS clusters to the opposite cell sides [Cha+20]. Respective forces pulling the CSs to the cell cortex are generated by cortical dynein. Note that dynein in fact promotes the CS clustering [Qui+05] rather than separation of the CSs, however, we reiterate that dynein has multiple functions in mitosis and is regulated differently in different parts of the cell. We hypothesize that both torque and dynein-mediated attraction of the CSs to the cell boundary are integrated to make the spindle bipolarity more robust.

The notions of torque and pivoting in MT-motor systems appeared in a few recent studies. Molecular motors generate torque when they move along the helical MT lattice [Nov+18]. Multiple motors crosslinking MT pairs can generate a bundling torque [Lam+19]. There may exist a torque between a CS and an MT anchored into the CS [End+94]. Torques of unknown origin in the spindle generate chiral MT structures [Mit+20]. Pivoting movements of MTs in spindles also have been observed [Kal+13; Win+19; FDA21]. The idea of the progressive restriction of the angle between a MT and a kinetochore to which this MT binds proposed in [Ede+20] is similar to our idea of the CS-CH torque. Such torque would most naturally emerge if MTs were cantilevered into kinetochores at normal angles, and if angular deformations of such connections were resisted elastically. However, there is little evidence of such elasticity of the MT-KTs connections. Indeed, plastic angular displacements of the kinetochores were observed instead [Lon+07] and swinging of MT bundles (K-fibers) projecting from the kinetochores over wide angles was observed [Sik+14; Elt+14]. Therefore, we hypothesize that the CS-CH torque could arise due to the geometric effect of the polar ejection force asymmetrically pushing on the proximal/distal parts of the CH arms, with the centromere pulled toward the CS, effectively rotating them into the orientation perpendicular to the CS-CH vector.

Our models have many limitations. To mention but two: (i) CH arms are very deformable, so modeling them as rods is not very accurate; (ii) our 2D solutions miss characteristic ‘doughnut-like’ spatial organization of the CH group in the spindle [Mag+15].

Although, prior studies have used non-local models for MT-motor-organelle organization [Man+18] or local PDE models to describe the spindle MT array [OJB20], our non-local description of spindle organization appears novel. The resulting model bares great similarity to descriptions of animal grouping [MEK99; LRC00; TB04; CK14] and models of particles interacting with orientations [MEK96; OHS17; OEK18]. Exploring analogies and lessons from these fields may create helpful intuition for understanding intracellular architecture or inspire new mathematical pursuits in further understanding possible behaviors of these models.

## Acknowledgements

CEM and AM are supported by the National Science Foundation grant DMS 1953430 to AM. The authors disclose no conflict of interests.

## S1 Table of parameter values

| parameter | meaning | value(s) in particle sim | value(s) in PDE sim |
| --- | --- | --- | --- |
| $f_a$ | strength of chrom-CS long-range attraction | 1 | 1 |
| $f_r$ | strength of chrom-CS short-range repulsion | 5 | 5 |
| $f_c$ | strength of chrom-chrom repulsion | 1.5 | 1 |
| $f_s$ | strength of CS-CS repulsion (+) or attraction (-) | varied between $-10, 2$ | varied between $-2, 2$ |
| $L_a$ | lengthscale of chrom-CS long-range attraction | 15 | 15 |
| $L_r$ | lengthscale of chrom-CS short-range repulsion | 5 | 5 |
| $L_c$ | lengthscale of chrom-chrom short-range repulsion | 1 | 1 |
| $L_s$ | lengthscale of CS-CS attraction or repulsion | 7.5 | 5 |
| $\tau_c$ | strength of chrom-chrom torque | 0.5 | 0.2 |
| $\tau_s$ | strength of chrom-CS torque | varied between 0, 5 | varied between 0.5, 1.5 |
| $D_C$ | chrom diffusion coefficient | 0 | 0.2 |
| $D_s$ | centrosome diffusion coefficient | 0 | 0.2 |
| $N_C$ | number of chromosomes | varied, 20 in most simulations | varied, 20 in most simulations |
| $N_S$ | number of centrosomes | varied, 10 in most simulations | varied, 10 in most simulations |

## S2 Annular equilibria of modified system

The results of the 1d stability analysis suggest that a monopolar spindle should evolve when CSs attract each other, so that all CSs concentrate in a small region of space, and the CHs tend to concentrate at a certain distance from the CSs, where the CS-CH centromere attraction balances the CS-CH arm repulsion. In 2D this will look like a ring pattern with CSs at the origin, and CHs in the ring with the center at the origin. Similarly, when CSs repel each other, one would expect that a ring of the CSs around the CH cluster in the center could be the steady state in the 2d continuous model. In this section, we verify these equilibria analytically in the 2d continuous model *without diffusion, alignment and torque terms*. To do so, we rely on the results of [FHK11; BLL12; CK14; Ber+15] that observe that for a certain choice of Newtonian interaction potentials between interacting particles, some analytical solutions in 2d interacting particles models can be found explicitly.

**Figure S1:**
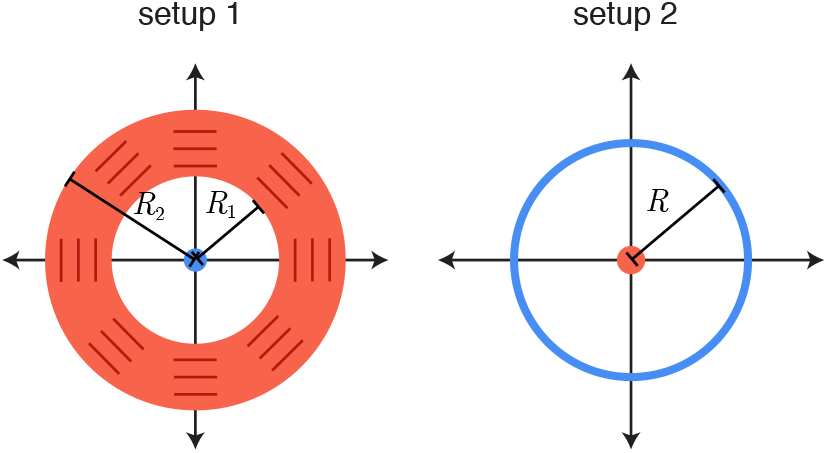
Two annular equilibria considered in the supplement. The explicit calculation is for a monopolar solution, shown in setup 1, but setup 2 (multipolar) follows from similar arguments.

In order to do so, we omit the diffusion and torque terms in the model, and also modify the distance dependent of the forces as follows:

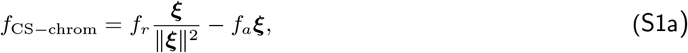

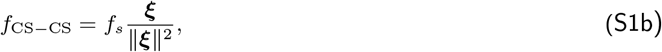

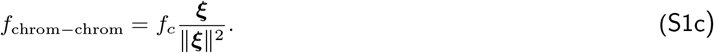

Here ***ξ*** is the 2d vector between interacting particles ***ξ*** = ***x*** − ***x***′. Note that in this case, the force amplitudes *f*_*r*_, *f*_*a*_, *f*_*s*_, *f*_*c*_ are the same as introduced above; however, the distance dependence of the forces is not exponential. Instead, the force distance dependence is scale-less, and the forces are the power functions. Namely, the attraction between the CS and the CH centromere is spring-like, proportional to the CS-CH mutual distance, while all three other forces decrease as the inverse distance, similar to the gravity and electrostatic forces.

The density equations, in the absence of the diffusion terms, become drift equations and can be solved by the method of lines. The characteristic curves for the CH density starting at *t* = 0 at position ***x***_0_ satisfy:

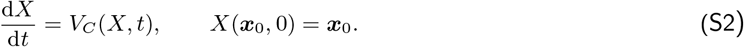

The CH density itself along these characteristics, *x* = *X*(*x*0, *t*), satisfies:

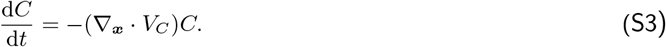

The velocity of the CHs is determined by (assuming *τ* = 0)

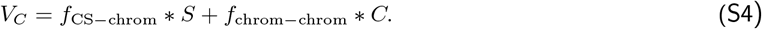

The analytical solutions are enabled by the fact that, in 2d only

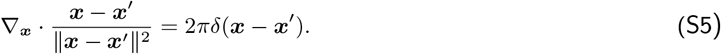

We assume that the monopolar solution is characterized by 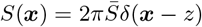. In this case

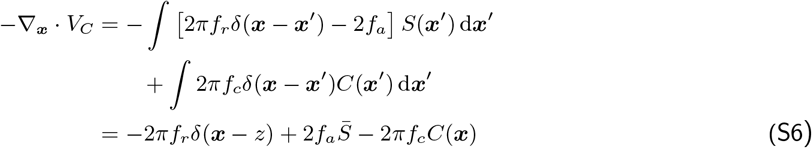

We look for the solution in which the CS and CH populations do no overlap, so *x* ≠ *z* and

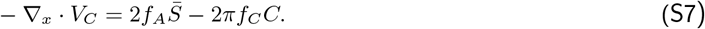

Plugging this expression into the characteristic equation, we have

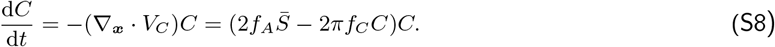

Since *C* > 0, this means that the CH stationary density is constant, and satisfies

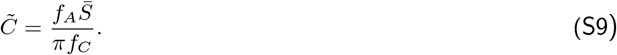

We now also resort to the observation that in 2d

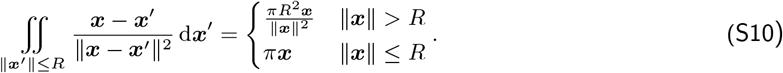

Let us consider the annular region, in which the CH density is non-zero: *A* = *R*_1_ ≤ ||***x***|| ≤ *R*_2_. We have

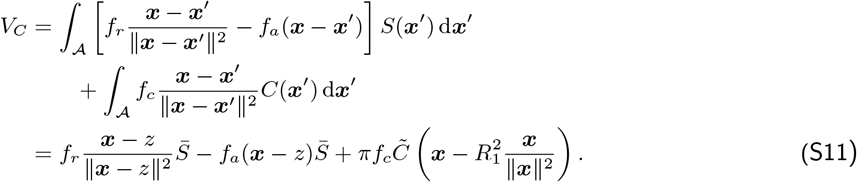

Assuming that the CSs aggregate at the origin (*z* = 0), so 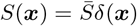, this reduces to:

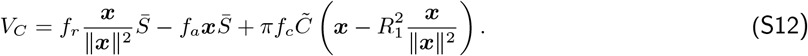

Since this velocity must be uniformly equal to zero, we have the following conditions:

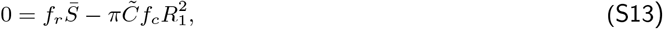

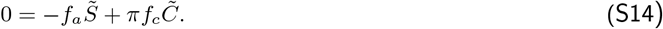

The first condition tells us that:

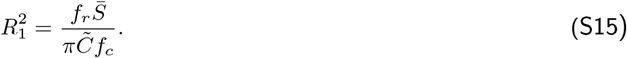

The second condition looks like a new condition, but is exactly equivalent to (S9) after algebra. The outer radius is then constrained by the conservation of the total CH number, manifesting as:

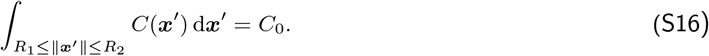

Consequently, this is indeed an equilibrium of the model.

A nearly identical calculation shows that the configuration with CHs concentrated at the origin and mutually repulsive (*f*_*s*_ > 0) CSs distributed along an annulus with the center at the origin, corresponding to the multipolar spindle, is also a valid solution of the 2d continuous model, if CH self-repulsion, torque and diffusion terms are neglected. Investigating the stability of each these ansatz solutions by using the perturbation technique of [CK14] seems natural, however, we want to understand the solutions in the presence of torque. We were unable to complete the stability analysis with torque seemingly because the torque and linear force terms are *not* separable as a product. We therefore resort to numerical solutions of the 2d continuous and discrete models. This is an interesting avenue of future study: whether stability analysis on the full model is possible, or if there is a viable approximation to the torque terms that enables this analysis.

## Notes

### Competing Interest Statement

The authors have declared no competing interest.

